# Effect of Dual Leucine Zipper Kinase Inhibitor GNE-3511 on Epileptogenesis and Cognitive and Behavioral Changes in a Temporal Lobe Epilepsy Model in Mice

**DOI:** 10.1101/2023.06.11.544443

**Authors:** Dilan Acar, Emre Tunçcan, Serra Nur Selek, İrem Ekin Sayın, Can Berk Asaroğlu, Gamze Tümentemur, Devrim Öz Arslan, Samet Özdemir, Güldal Güleç Süyen

## Abstract

Inflammation and neuronal loss are key factors in the pathophysiology of epilepsy. It is known that activation of Dual Leucine Zipper Kinase (DLK) in chronic neurodegenerative diseases causes neuron death and axon degeneration through apoptotic pathways. Genetic deletion of DLK (dual leucine zipper kinase, MAP3K12) or pharmacological inhibition by GNE-3511 or GNE-8505 was found to have benefits in mouse models of Alzheimer’s and ALS; however, there is no previous study that uses DLK inhibitors in epilepsy models. Therefore, the aim of our study is to suppress the DLK/JNK pathway with the DLK inhibitor GNE-3511 in the temporal lobe epilepsy model induced by pilocarpine and to determine the effect of this inhibition on cognitive and behavioural defects, as well as epileptogenesis. Open field, elevated plus maze, and Morris water maze tests were used to assess locomotor activity, anxiety and learning and memory, respectively. Following decapitation the hippocampi were removed and, histopathological and biochemical analyses were performed. Both treatment groups had less anxiety and locomotor activity as well as decreased impairment in memory and learning performance compared to SE group (p<0.001). Both doses of GNE-3511 prevented the spontaneous recurrent seizures in a dose dependent manner with respect to the SE group (p<0.01 and p<0.001, respectively). Histological examination of the hippocampus revealed a neuroprotective effect of both doses of treatment in the CA1 area and dentate gyrus (p<0.0001). Thus, DLK inhibitor GNE-3511 was found to be effective in preventing epileptogenesis, neuronal loss and cognitive and behavioural deficits due to pilocarpine induced SE.

## 1. Introduction

Epilepsy is a heterogeneous brain disease that has a persistent tendency to produce epileptic seizures and is characterized by neuro-biological, cognitive, psychological and social consequences (***Fisher et al., 2014***). Although the cause of epilepsy is mostly unknown; all kinds of factors that may affect the function of the brain, traumatic brain injury, infection and genetic mutations, can cause epilepsy (***Dooling and Costa-Mattioli, 2018***). The mainstay of treatment is antiepileptic drugs, and these drugs act by suppressing seizure formation without correcting the underlying neuropathological process of the disease. However, approximately 30% of epilepsy patients show resistance to these conventional antiepileptic drugs (***Chen et al., 2017***). This situation reveals the need to develop new treatments and to identify epilepsy/epileptogenesis biomarkers (***Caro et al., 2019***).

The process of epileptogenesis includes the loss of the balance between excitatory and inhibitory activity in a neuronal network, resulting in disruption of neuronal processes and neuronal losses (*Fisher et al., 2005*). It is thought that the initial brain injury results in hippocampal cell loss, and then collateral axonal sprouting and reorganization of synaptic circuits affect the balance between inhibition and excitation in limbic circuits until spontaneous seizures finally occur (***Jensen et al., 2012; Thijs et al., 2019***).

Neurocognitive disorders are frequently observed in epilepsy patients. Mental slowness, memory difficulties and attention deficits constitute most of the cognitive complaints in adult patients (***Rijckevorsel, 2006***). Temporal lobe epilepsy (TLE) is the most common type of refractory focal epilepsy in adults. Despite the increase in antiepileptic drugs developed in recent years, it is known that 30% of TLE patients are drug-resistant (***Schuele and Lüders, 2008***). The experimental TLE model created with pilocarpine in experimental animals is a valid model that is frequently used in research (***Leite et al., 1990***)

Studies on the pathogenesis of epilepsy have shown that cell death due to epileptic seizures can occur through the apoptotic process. While the mechanism of epileptogenesis is still hypothetical, there is a growing body of studies showing that inflammation also plays a prominent role. While acute brain injuries such as traumatic brain injury, central nervous system (CNS) and stroke may cause epilepsy, inflammation and neuronal loss are key factors in the phenomenon (***Vezzani et al., 2011; Klein et al., 2017***).

c-Jun N-terminal kinases (JNKs), members of the mitogen-activated protein kinase (MAPK) family, another important pathway triggered by extracellular stress in epilepsy, are activated in response to a wide variety of stimuli. It is particularly important that the cell is exposed to various biotic or abiotic stress events such as inflammation, oxidative stress, deoxyribonucleic acid (DNA) damage, osmotic stress or cytoskeletal changes (***Urano et al., 2000***). c-Jun-N-terminal kinases also regulate important physiological processes such as neuronal functions, immunological effects, and embryonic development through their effects on gene expression, cytoskeletal protein dynamics, and cell death/survival pathways (***Zeke et al., 2016***). In a study conducted with the pilocarpine model, aggregation and hyperactivation of JNK isoforms in the CA1 region of the hippocampus of chronic epileptic rats with frequent convulsive seizures was demonstrated (***Tai et al., 2017***).

Stress-specific JNK activation occurs with DLK activation in neurons via the MKK4/7 pathway, and this activation increases PERK signalling by a yet unknown mechanism (***Ghosh et al., 2011; Simon et al., 2016; Larhammar et al., 2017***). Induction of these pathways leads to widespread transcriptional damage through regulation of transcription factors such as c-Jun and adenosine triphosphate, which lead to apoptosis and axon degeneration in many neurons. In chronic neurodegenerative diseases, neuron death and axon degeneration occur with the activation of the stress pathway related to DLK (***Simon et al., 2016; Watkins et al., 2013; Shin et al., 2012***).

In a study using genetically engineered mice in which the dual leucine zipper kinase gene was deleted, it was shown that the degeneration of neurons after axotomy or vincristine treatment is dependent on DLK (***Simon et al., 2016; Miller et al., 2009***). Similarly, in the in vivo optic nerve damage model, neurons without DLK expression were found to be protected from neurodegeneration (***Watkins et al., 2013***). In other study, it was shown that loss of DLK activity protects neurons from excitotoxicity-induced degeneration in a kainic acid-induced excitotoxicity model in DLK knockout mice. Mice in which the DLK gene was deleted in all tissues as part of the same study were examined 3 months after DLK deletion and found no histological, electrophysiological, or behavioral abnormalities (***Pozniak et al., 2013***). This suggests that loss of DLK signal following inhibitor therapy will be well tolerated.

To our knowledge, there is no previous study using DLK inhibitors in epilepsy models. The aim of the present study is to determine the effect of suppression of the DLK/JNK pathway by the DLK inhibitor GNE-3511 on epileptogenesis and cognitive and behavioural deficits in pilocarpine induced temporal lobe epilepsy model in mice.

## 2. Results

### 2.1 Monitorization

The number of spontaneous seizures per day decreased significantly in both the 1 mg/kg (p<0.01) group and the 5 mg/kg (p<0.001) group with respect to the SE group. There was no significant difference in the spontaneous seizure scores between groups.

### 2.2 Cognitive and Behavioral Assessments

#### 2.2.1 Open Field

SE group had significantly increased number of squares visited with respect to the control group (*p<0.001*). Both doses of GNE-3511 prevented the increase in locomotor activity (*p<0.01*). (Fig. 2).

**Fig. 1.**
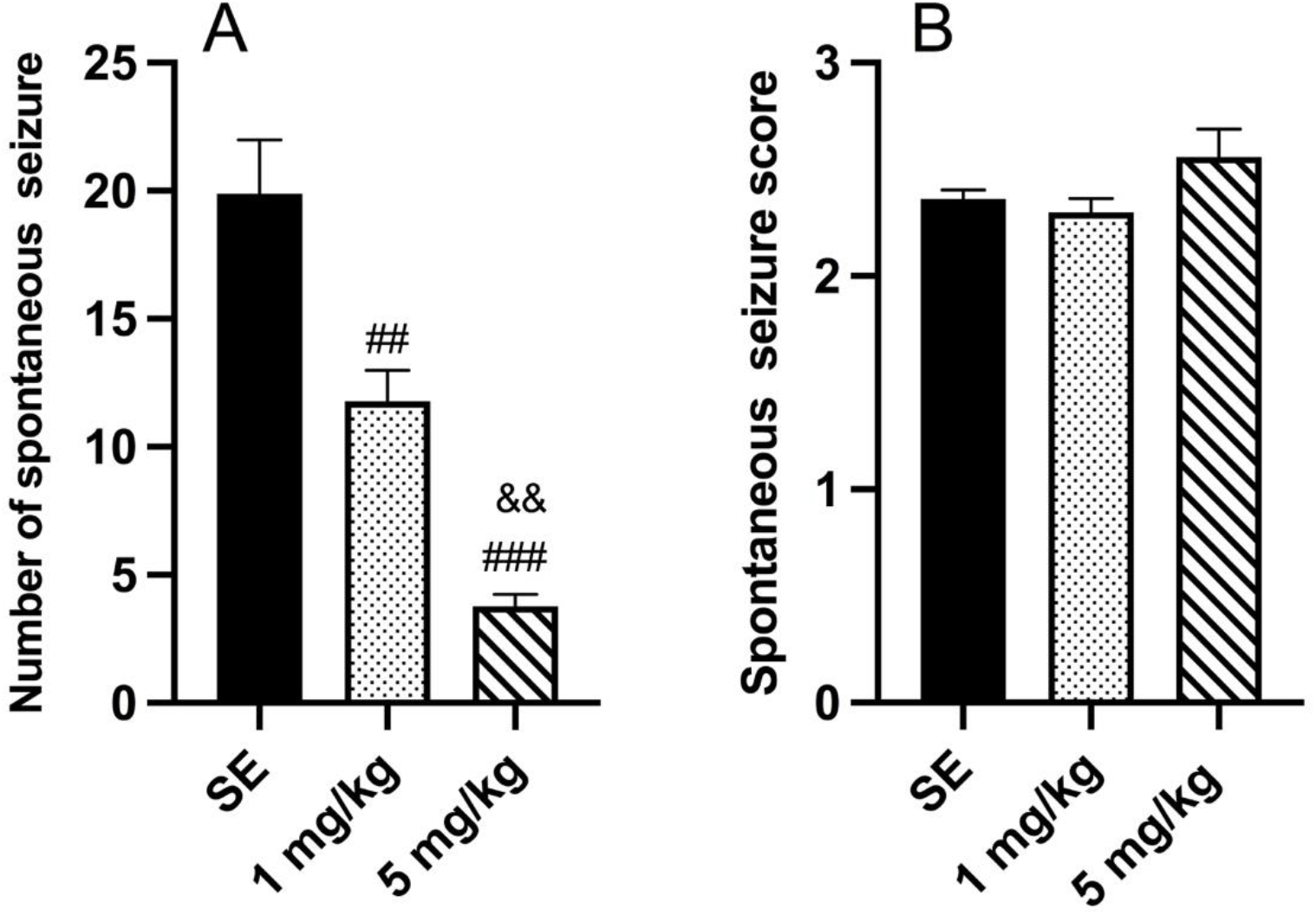
Daily spontaneous seizure numbers (A) Spontaneous seizure scores (B). Results were presented as means ± S.E.M. Each group consisted of 6-8 mice. ^##^*p<0.01*, ^###^*p<0.01* vs SE group and ^&&^*p<0.01 vs 1 mg/kg GNE-3511 group*

**Fig. 2.**
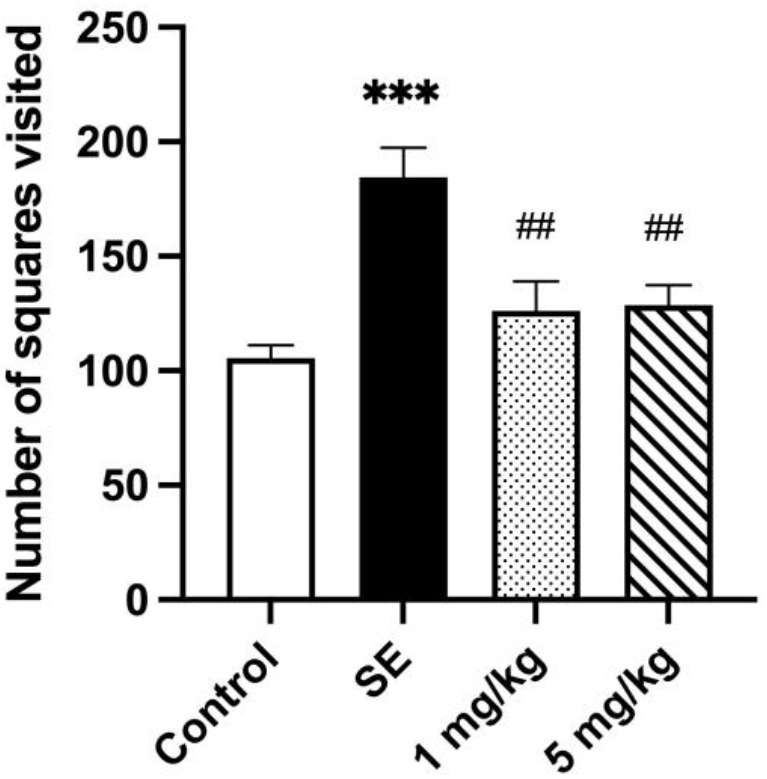
Locomotor activity was assessed by the open field test. The number of squares visited was recorded. Results were presented as means ± S.E.M. Each group consisted of 6-8 mice. ^***^*p<0.001* vs control group. ^###^*p<0.01* vs SE group.

#### 2.2.2 Elevated Plus Maze

The SE group mice spent significantly less time in the open arms compared to the control group mice (*p<0.001*). The anxiety score was significantly lower in the SE group compared to the control group (*p<0.0001*). Both 1 mg/kg and 5 mg/kg doses of GNE-3511 prevented the decrease in anxiety scores due to SE (*p<0.0001*) (Fig. 3C).

**Fig 3.**
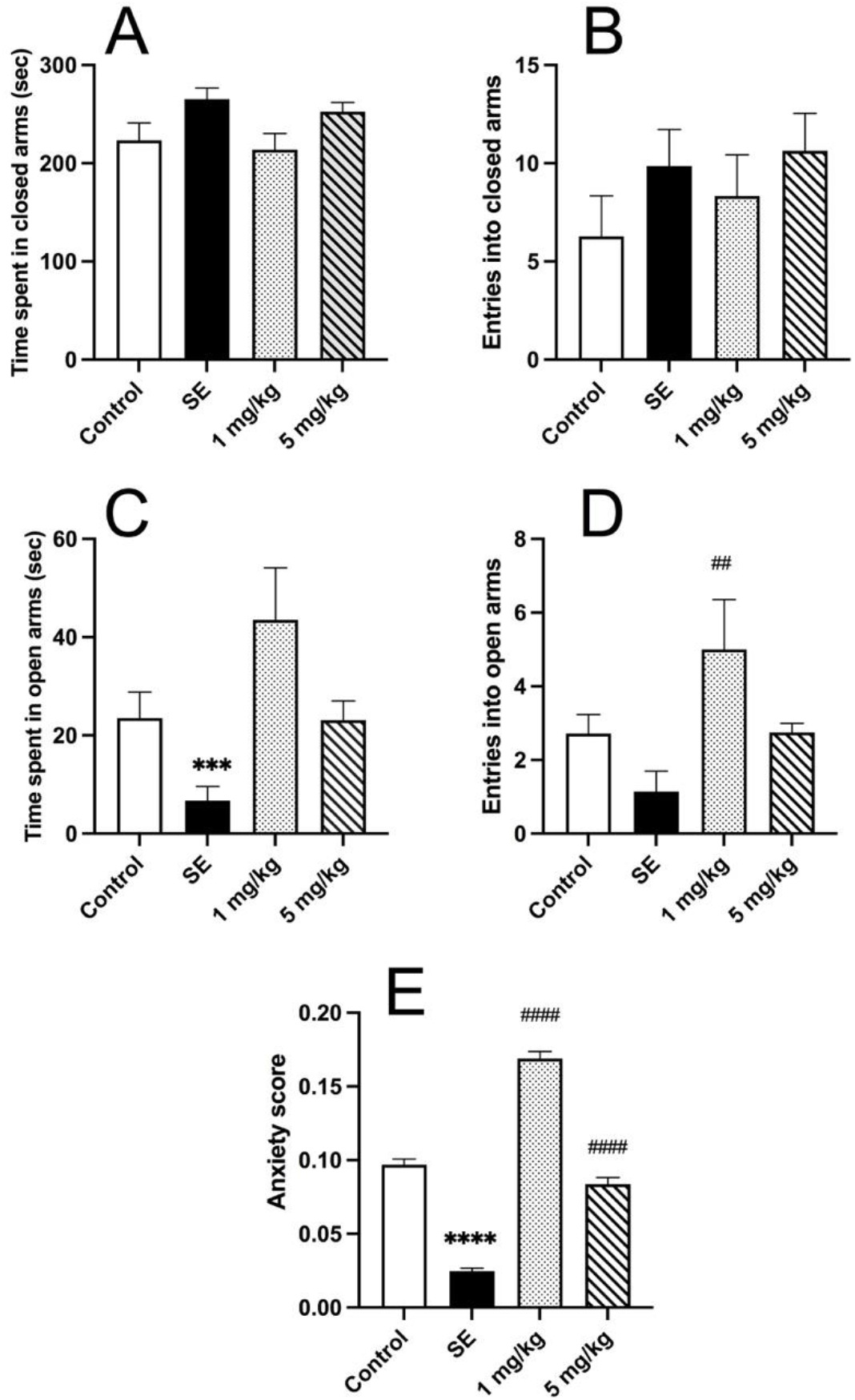
Anxiety behavior was assessed on the elevated plus maze test. Time spent in open arms (C) and anxiety scores were calculated (time spent in open arm/ time spent in open arm+time spent in closed arm) (E). Results were presented as means ± S.E.M. Each group consisted of 6-8 mice. ^****^*p<0.0001*, ^***^*p<0.05* vs control group, ^####^*p<0.0001*, ^##^*p<0.01* vs SE group.

#### 2.2.3 Morris Water Maze

The daily escape latency data is shown Fig. 4. Escape latency decreased significantly in the control group, and 1 mg/kg and 5 mg/kg treatment groups (*p<0.001*) compared to the 1st training day. On the other hand, no significance was found in the SE group in terms of acquisition.

**Fig. 4.**
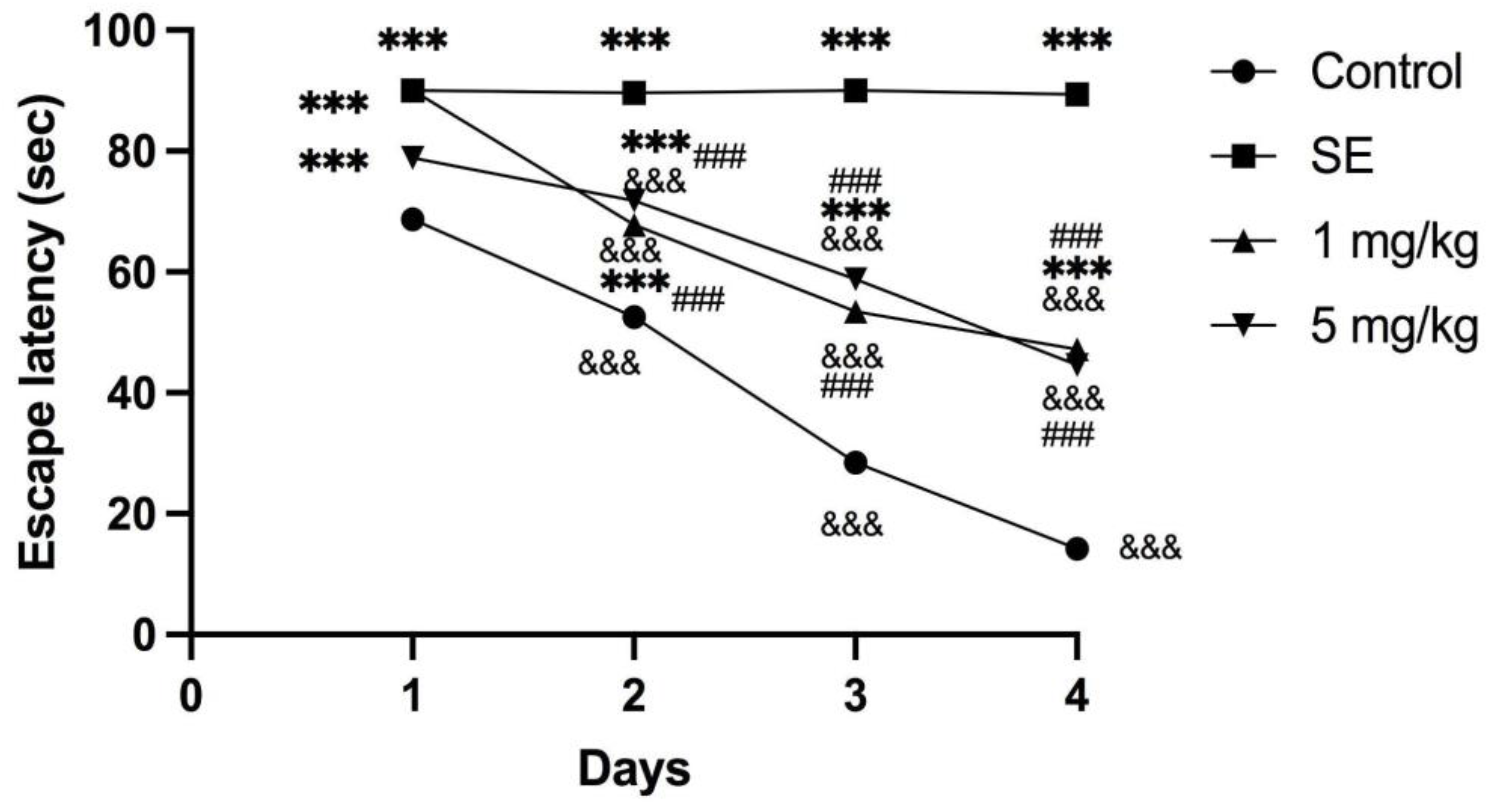
Morris water maze four training days of acquisition performance. Results were presented as means ± S.E.M. Each group consisted of 6-8 mice. ^***^*p<0.001* vs control group, ^###^*p<0.001* vs SE group and ^&&&^*p<0.001* with respect to their first performance day.

Comparison of the escape latency on a daily basis revealed a significant difference between the control and SE groups (*p<0.001*). Both 1 mg/kg and 5 mg/kg treatment groups spent less time finding the platform than the SE group (*p<0.001*) but spent more time compared to the control group (*p<0.001*). There was no significant difference between 1 mg/kg and 5 mg/kg treatment groups (Fig. 4A).

When the time spent in the target quadrant on probe day was analysed there was no significance between treatment groups (Fig. 5B). On the other hand, the number of platform crossings decreased significantly in the SE group with respect to the control group (*p<0.05*) (Fig. 5A).

**Fig. 5.**
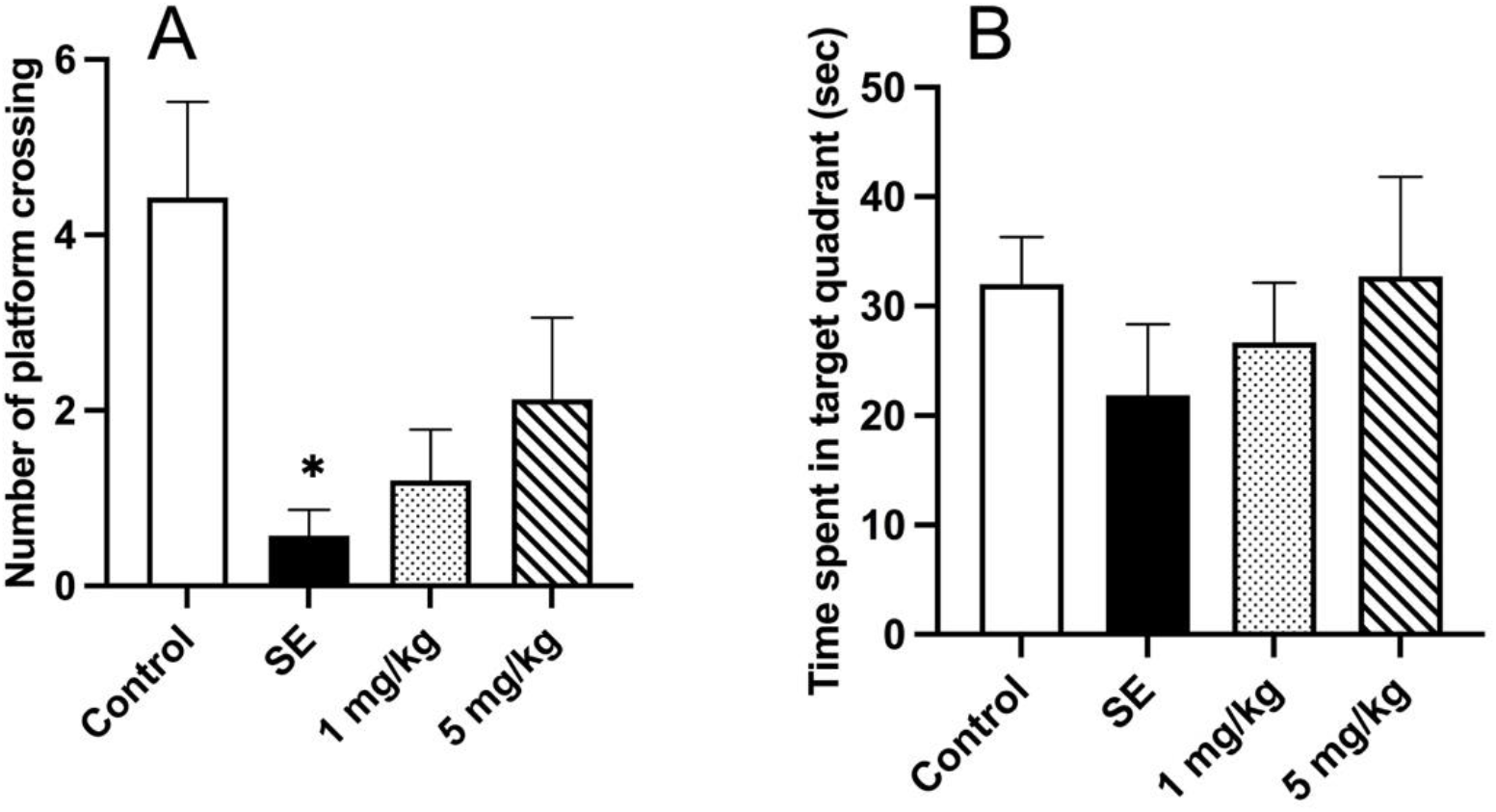
Morris water maze retention performance. Results were presented as means ± S.E.M. Each group consisted of 6-8 mice. (A) Number of platform crossings and (B) time spent in the target quadrant were calculated. ^*^*p<0.05* vs control group.

### 2.3 Histopathological analyses

The histological examination of H&E-stained sections from the hippocampus revealed necrosis and degeneration in the SE group (Fig. 6B). The cell bodies of CA1 neurons in the 1 mg/kg GNE-3511 treated group were arranged regularly and the condensed nuclei disappeared (Fig. 6C). In the 5 mg/kg GNE-3511group, less irregular neurons was shown in cells of the pyramidal layer of the CA1 area (Fig. 6D).

**Fig. 6.**
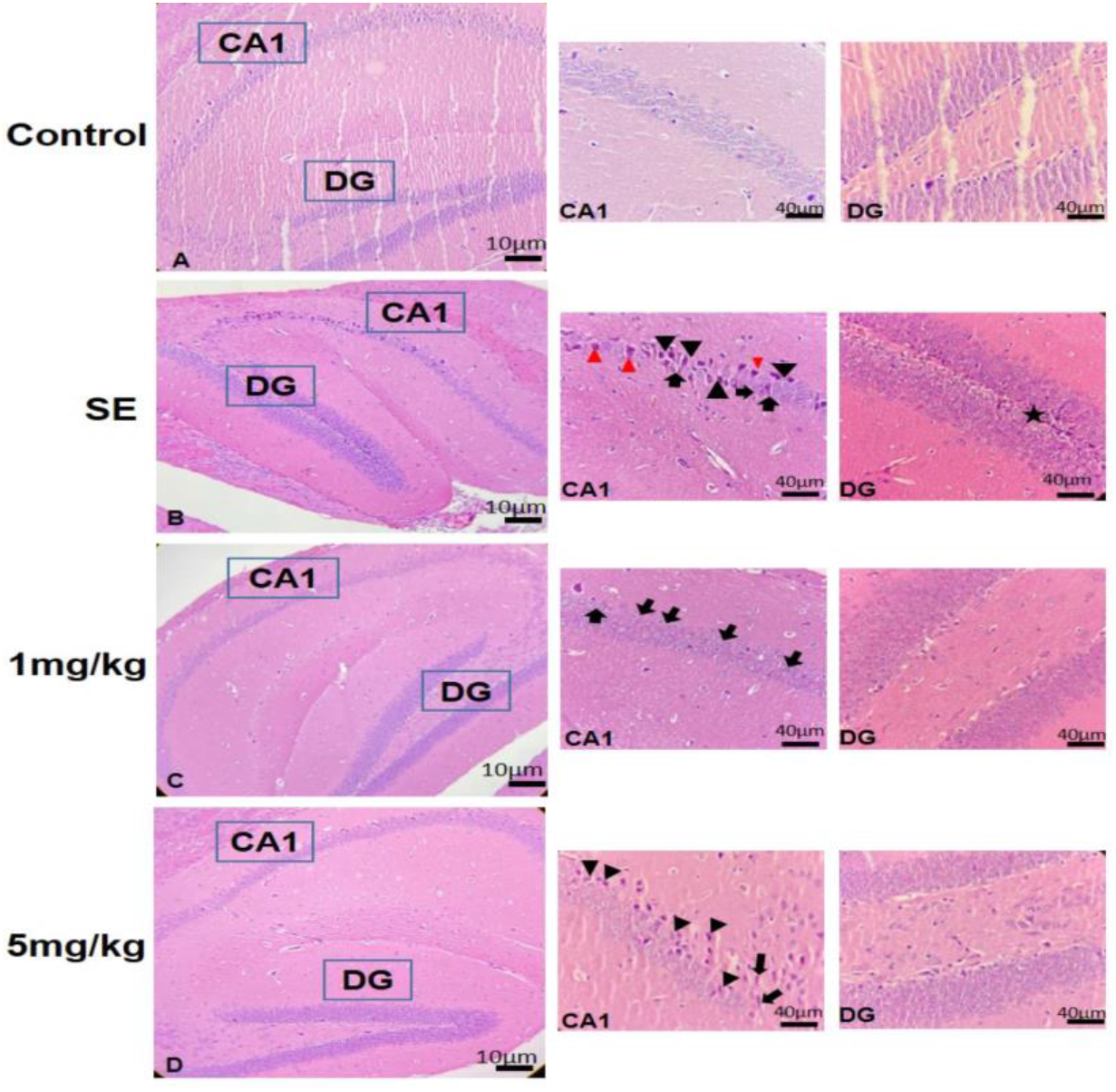
The morphological changes of the hippocampal CA1/DG region. A) The control mice show normal histological structures of CA1 and DG regions. B) The hippocampus of the SE group shows extensive damage in the CA1 and dentate gyrus cells. Black arrows indicate degeneration of neurons, black arrowheads head indicate necrosis of neurons, red arrowheads indicate nuclear deviation in CA1, and star indicates mostly marked pyknotic neurons in DG. C) The boundaries between cytoplasm and nucleus are clear in the 1 mg/kg GNE-3511 group. Many intact and round cells (black arrows), neurons closely arranged in DG. D) Arrow indicates less irregular neurons in CA1, arrowheads indicate microglia in DG of the 5 mg/kg GNE-3511 group.

In the SE group, degeneration and necrosis of neurons accompanied by a significantly increased number of dead neurons were observed in the CA1 and DG regions compared with the control group (p<0.0001). In contrast, both doses of GNE-3511 treatments caused neuroprotection in the CA1 area DG (Fig. 7).

**Fig. 7.**
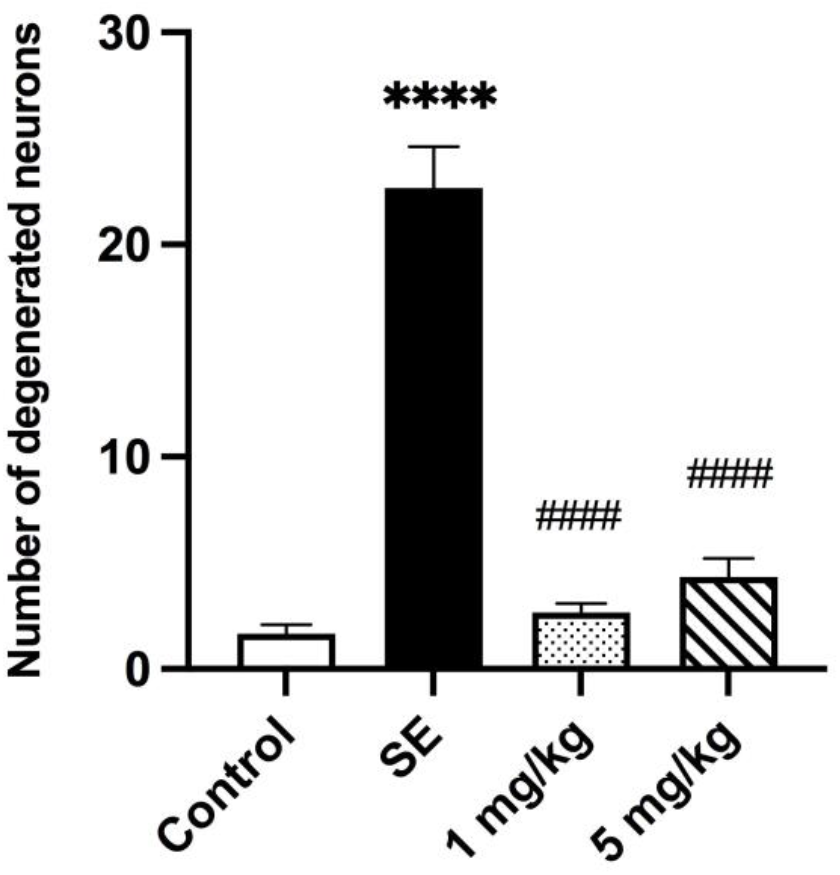
Number of degenerated neurons. The final value presented as means ± S.E.M. ^****^*p<0.0001* vs control group, ^###^*p<0.001* vs SE group.

### 2.4. Western blot

The c-Jun protein levels were evaluated in all groups. SE group had significantly higher levels compared to control (*p<0.05*), 1 mg/kg (*p<0.0001*), and 5 mg/kg treatment groups (*p<0.0001*). Treatment with GNE-3511 decreased c-Jun levels in SE mice (Fig. 8).

**Fig. 8.**
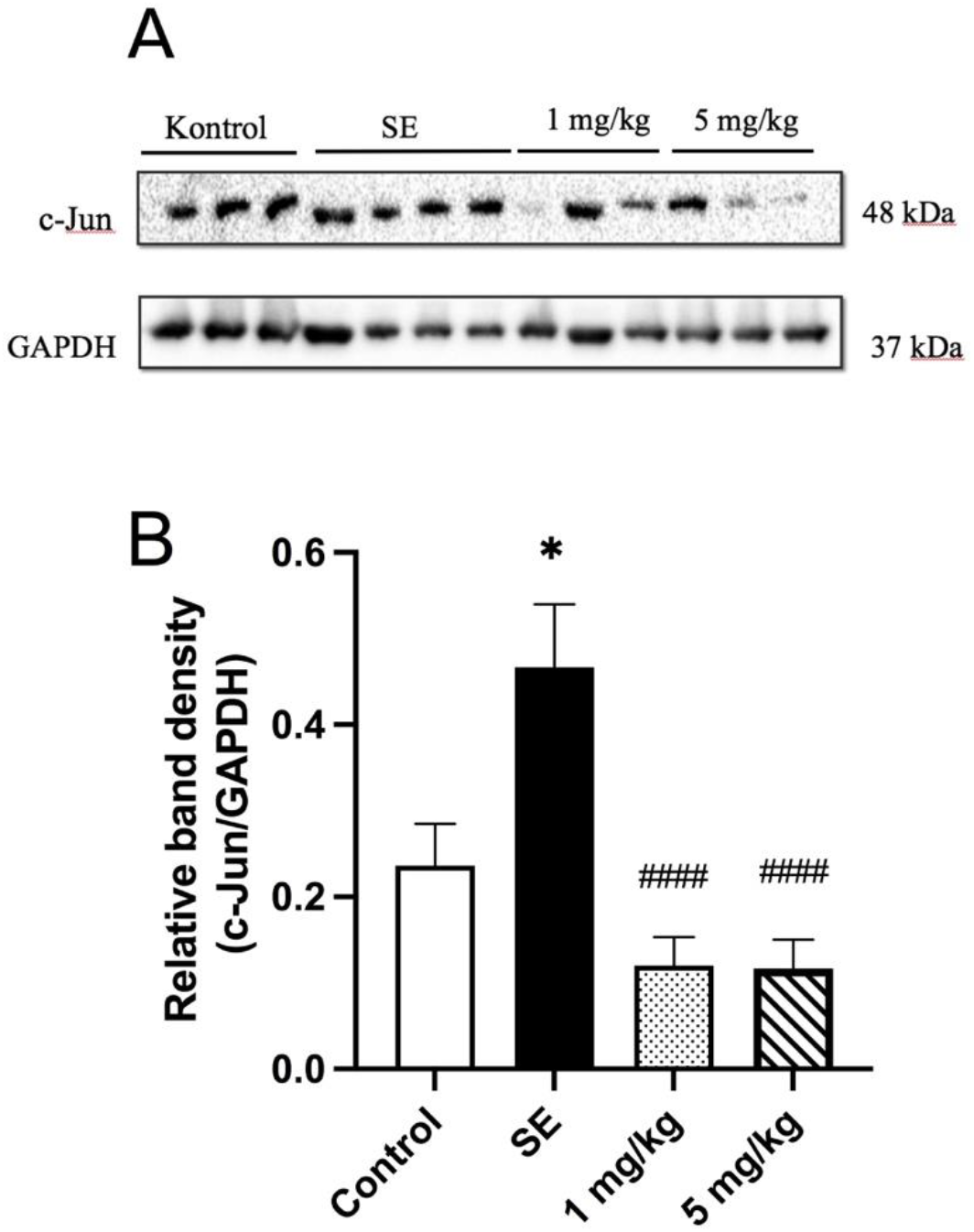
Representative western blot analysis of c-Jun protein levels in hippocampal tissue. c-Jun protein bands (A) were quantified by ImageLab software. GAPDH was used as an internal loading control. Relative band density (c-Jun/GAPDH) (B). The final value presented as means ± S.E.M of at least 3 biological replicates. ^*^*p<0.001* vs control group, ^####^*p<0.001* vs SE group.

## 3. Discussion

Epilepsy is a common neurological disease characterised by recurrent seizures. Temporal lobe epilepsy (TLE) is one of the most common forms of focal epilepsy in adults (***Begley et al., 2015***). It is known that DLK plays an important role in acute neuronal injury and chronic neurodegenerative diseases (***Siu et al., 2018***). There is no study in the literature in which DLK inhibitors were tested on the TLE model. For this reason, the effects of two doses of DLK inhibitor GNE-3511, 1 mg/kg and 5 mg/kg, on epileptogenesis, cognitive and behavioural disorders as well as neuronal damage, were investigated in this study.

In our study, we used the pilocarpin model to mimic the epileptogenesis and to understand the effect of the DLK inhibitor GNE-3511 on seizures. Pilocarpine initiates seizures with the activation of the cholinergic system, namely M1 muscarinic receptors, and activates the NMDA (N-methyl-D-aspartate) receptor for seizure maintenance (***Hamilton et al., 1997***). SE, 1 mg/kg and 5 mg/kg groups, when seizure activity was followed for 8 hours a day for 10 days, we found that GNE-3511 reduced the number of seizures, especially in the high dose group. The JNK pathway has emerged as an important regulator of neuronal degeneration induced by NMDA receptor hyperactivation. Studies have shown that inhibitors of JNK activity can prevent NMDA-mediated neuronal apoptosis both in vitro and in vivo after ischemia (***Borsello et al., 2003; Gao et al., 2005***). Considering the results, it is suggested that DLK may reduce seizures through JNK-mediated NMDA receptor regulation.

Open-field test is used to analyse locomotor activity, anxiety, and stereotypical behaviours (***Prut et al., 2003***). Changes in locomotor activity may be indicative of altered neurological processes. Therefore, it may reflect abnormal brain functions. The fact that the experimental animal spends more time in the open area may indicate that the stress level is lower (***Kraeuter et al., 2019***). In addition, in some epilepsy studies, it was observed that motor activity increased in SE-induced animals (***Stewart et al., 2003***; ***Szyndler et al., 2005***). In our study, locomotor activity was analysed in the open field test and it was observed that locomotor activity in the SE group significantly increased compared to the control and treatment groups. At the same time, the locomotor activity of the 1 mg/kg and 5 mg/kg treatment groups was higher than the control group. According to the results, it was observed that locomotor activity in the treated groups approached the control group. It has been reported that the increased activity in pilocarpine-induced SE models may result from abnormal activity of the hippocampal-accumbens pathway that affects locomotion (***Stewart et al., 2003***; ***Groenewegen et al., 1999***). Hippocampal afferent neurons in the nucleus accumbens interact with dopaminergic fibers from the ventral tegmental area, forming an effective functional interface between the limbic and motor systems. In one study, electrical stimulation of the hippocampus or local injections of glutamate receptor agonists in the nucleus accumbens were found to increase locomotor activity in rats (***Donzanti and Uretsky, 1983***). In another study, both interictal activity in the CA1 region of the hippocampus and increased locomotor activity were observed in the same period following pilocarpine induction. However, changes in this activity pattern were seen without cell loss in the suprachiasmatic nucleus or intergeniculate leaf. Therefore, it has been suggested that the increase in locomotor activity may be due to functional changes caused by seizure activity in these regions (***Stewart et al., 2003***). In this context, the findings in our study that the increased locomotor activity following SE decreased after DLK inhibition suggests that it may have affected the seizure activity with a different mechanism.

Anxiety is a negative emotional state associated with the perception of threat. It is characterised by the expectation of potential threats or negative consequences. A step towards a better understanding of the pathophysiology of this mood is the use of animal models (***Sarnyai et al., 1987***). The elevated plus maze test is based on the tendency of the less anxious animal to visit the open arm of the maze more, while the more anxious animal to spend more time in the closed arm (***File, 1987***). While some studies in epilepsy showed that anxiety-like behaviour increased (***Vrinda et al., 2017***), it was decreased in some (***Gulec et al., 2016***). Nevertheless, increased anxiety in epilepsy has been reported both in reports of TLE patients and in pilocarpine-induced TLE models (***William et al., 2016***). In our study, SE group spent less time in the open arm compared to the control and treatment groups. In addition, 1 mg/kg treatment group spent more time in the open arm than the control group. Experimental evidence reveals that chronic epilepsy causes extensive damage to the ventral and dorsal hippocampus and amygdala, especially in the pilocarpine-induced epilepsy model (***Inostroza et al., 2012***). It is thought that differences in the size of the amygdala lesion may be responsible for the differences in threat perception (***Inostroza et al., 2011***). These observations suggested that differences in behavioural reactivity may be related to underlying brain injury rather than the epileptic state (***Inostroza et al., 2012***). Therefore, it can be considered that DLK inhibition has a neuronal protective effect on the targeted brain injury and a reducing effect on the increased anxiety associated with SE.

Memory disorders and complaints are frequently encountered in epilepsy. The majority of these complaints are composed of TLE patients whose seizures are directly related to memory-related brain structures (***Kapur and Prevet, 2003***). Since the temporal lobe is the main structure that includes learning and memory, damage to these structures as a result of spontaneous seizures may cause some memory disorders (***Gilliam et al., 2003; Vrinda et al., 2018***). Morris water tank test was used to evaluate spatial learning and memory. In the evaluation of the 4-day performances of the groups within themselves, it was observed that the learning curve of the SE group did not change, and there was no significant decrease in the latency of finding the platform compared to the 1st day. In the control, 1 mg/kg and 5 mg/kg groups, the latency of finding the platform was significantly reduced compared to the first day, and the learning was achieved. Together with previous studies (***Detour et al., 2005; Pearson et al., 2014***) showing spatial learning and memory impairment in temporal lobe epilepsy, DLK inhibition with a GNE-3511 inhibitor had a curative effect on spatial learning.

When the differences between the groups in the platform finding on a day basis are examined, it is seen that the latency of 1 mg/kg, 5 mg/kg and control groups decreased significantly compared to the SE on the 2nd, 3rd and 4th days. When the number of platform crossings was evaluated, in the first 4 days of the 5th day probe trial was evaluated, it was observed that the crossing number of the SE group was lower than the control group. Other studies have shown memory impairments in SE models created with pilocarpine (***Detour et al., 2005; Pearson et al., 2014***).

Neuronal mechanisms related to SE-induced learning and memory disorders have not yet been elucidated. However, it is thought that permanent changes in long-term potentiation (LTP) and synaptic activity may be related to this disorder in the cellular context. In a study conducted with the pilocarpine-induced SE model, it was shown that LTP formation was impaired for 1 week by SE induction. It was thought that this situation might be related to the saturation of the synaptic response caused by epileptic activity (***Kryukov et al., 2016***). In another study, it was found that epileptiform activity leads to an increase in α-amino-3-hydroxy-5-methyl-4-isoxazolepropionic acid (AMPA) receptor-mediated CA3-CA1 synapse conduction due to N-methyl-D-aspartate (NMDA) receptors. Henceforth, it has been suggested that it inhibits additional synaptic potentiation (***Abegg et al., 2003***). In a study with GLuN2B antagonist, which is a subtype of NMDA receptor, it was observed that the decrease in LTP after SE was inhibited. Extrasynaptic GluN2B NMDA receptors have also been associated with neurotoxicity, degeneration and cell death. It is known that the DLK pathway is activated by extracellular stress factors and is associated with cell death (***Kryukov et al., 2016***). In our study, learning and memory performance, which is known to be impaired in temporal lobe epilepsy, improved with the DLK inhibitor GNE-3511. Therefore, the neuronal protective effect expected by DLK inhibition suggests that this mechanism may prevent learning and memory impairment after SE.

Hippocampal sclerosis is recognized as one of the main causes of focal epilepsy. It is also present in approximately 10% of adults with new-onset focal epilepsy. It is characterised not only by neuronal death, but also by changes in neuronal connectivity and network behaviour that underlie chronic epilepsy and memory disorders (***Walker, 2015***). Information on the mechanisms underlying hippocampal damage following prolonged seizures has largely been derived from animal models of SE (***Meldrum, 1991***). Neuronal damage results from excitotoxicity, which mediates neuronal death through activation of glutamate receptors during epileptic activity. In the present study neuronal damage in the CA1 area and DG induced by SE was prevented by both doses of GNE-3511 treatment. DLK is a mixed lineage kinase required for stress-induced JNK activity in neurons during development and is required for both neurodegeneration and axon regeneration of the dorsal root ganglion (***Ghosh et al., 2011***). The location of the DLK protein in the axon has led to the hypothesis that it acts as a damage sensor against axonal injury (***Watkins et al., 2013; Shin et al., 2012***). In addition, as a result of studies, it has been understood that DLK function is not limited to axonal damage, but is also necessary for neuronal damage response (***Pozniak et al., 2013***). The fact that DLK regulates the stress-induced JNK signal and that this signal makes a specific connection, unlike the JNK pathways in CNS diseases, means targeting neuronal dysfunction directly. Tolerating the loss of DLK in adults and the ability of a DLK inhibitor to reverse the transcriptional stress response induced by DLK signalling in the cell indicate a therapeutic potential (***Pichon et al., 2017***).

The JNK pathway causes activation of a number of transcription factors, including c-Jun, which are activated by stress stimuli. This activation causes various cellular responses such as apoptosis, cell differentiation and inflammation depending on the type and length of the stimulus (***Ferraris et al., 2013***). It has been shown that, DLK mediates neuronal apoptosis through the JNK pathway and c-Jun (***Chen et al., 2008***). In this study, Western blot method was used to measure the effect of the GNE-3511 on c-Jun protein levels. Relative band density analysis was performed for protein level determination. As a result, it was observed that the c-Jun protein level of the SE group was higher than both control and treatment groups. Decreased c-Jun levels in the treatment groups indicate that DLK inhibition is still effective at the protein level in the chronic phase of the TLE model.

In conclusion, we state that DLK inhibitor GNE-3511 prevented epileptogenesis and neuronal damage and improved the cognitive and behavioural changes induced by SE in the pilocarpine model of temporal lobe epilepsiy in mice.

## 4. Material and methods

### 4.1 Animals

All protocols were approved by the Ethic Committee of Acıbadem Mehmet Ali Aydınlar University (ACU-HADYEK 2021/07) and all treatments performed according to the guidelines of National Institutes of Health (NIH) for the Care and Use of Laboratory Animals.

Female C57BL/6 (18-25 g; n=53) mice were obtained from Acibadem Mehmet Ali Aydinlar University, Experimental Animals Research Center (ACUDEHAM). During all the course of the experiment, mice were kept under 12h/12 h light/dark cycles (lights on at 07.00 am), with appropriate humidity and temperature (22-25 ºC) levels. They were fed with pellet food and tap water ad libitum. Their cages were cleaned within 2-day periods.

Sample size was calculated at the beginning of the study using G*power statistical software, to achieve 95 % power at a significance criterion of α = .05

Animals were randomly divided into 4 groups:

Control group (n=7): No treatment

SE group (n=7): Induction of SE by pilocarpine followed by treatment with vehicle 20 ml/kg (%2 Tween80, %2 Glycerine, and double distilled water) (i.p. twice a day, 12 h interval) 1 mg/kg GNE-3511group (n=6) Induction of SE by pilocarpine followed by treatment for 7 days (i.p. twice a day, 12 h interval), 5 mg/kg GNE-3511 group (n=8) status epilepticus induced and treated for 7 days (i.p. twice a day, 12 h interval).

**Scheme 1.**
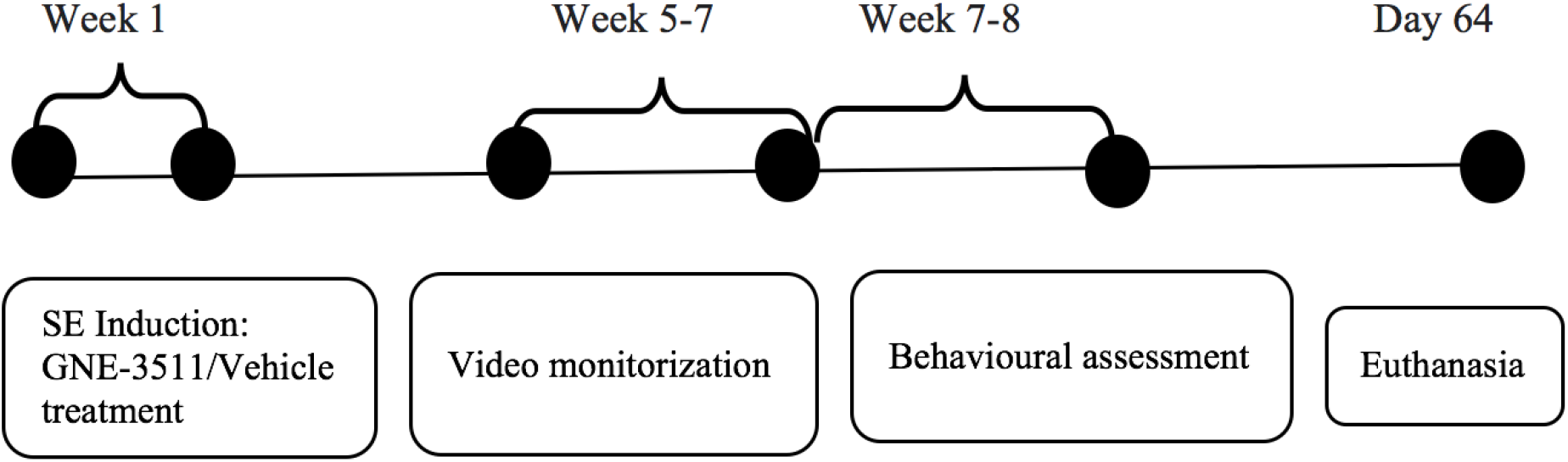
Study protocol

### 4.2 Induction of SE

Epilepsy was induced by pilocarpine. Animals were intraperitoneally (i.p.) injected scopolamine methyl bromide 30 min prior to pilocarpine injection in order to reduce its peripheral cholinergic effects. 100 mg/kg of pilocarpine was administered i.p. in every 20 min periods (maximum 7 doses). Ninety minutes after the onset of status epilepticus, seizures were suppressed with i.p. administration of 10 mg/kg diazepam to prevent the high mortality rate. GNE-3511 or vehicle treatment started 90 minutes after diazepam injection. Status epilepticus was determined using the Racine’s scale (Arshad and Naegele, 2020). Fifteen minutes after (GNE-3511 or vehicle) treatment, saline (1ml) was injected to all animals to reduce mortality and was repeated for 7 days (once a day). In addition, all animals were fed with baby food and injected 10% dextrose subcutaneously until they reached their initial weight.

### 4.3 GNE-3511 treatment

Mice were injected with 1 mg/kg or 5 mg/kg GNE-3511 i.p., dissolved in a 20 ml/kg solution for 7 days with 12 h intervals. GNE-3511 was formulated in a solution of %2 Tween80 v/v, %2 Glycerine v/v and double distilled water (ddH2O). The solution was sonicated and stored at -20 no more than 7 days.

### 4.4 Video monitorization of seizures

Four weeks after the end of the treatment, all mice except the control group were video-monitorized for 8 hours/day for 10 days to detect the frequency and duration of spontaneous recurrent seizures. To eliminate bias, each mouse was encoded with a random number before video monitorization and were evaluated blindly as well.

### 4.5 Cognitive and Behavioural Assessments

Open field test, elevated plus-maze test, and Morris water maze test were conducted in the Laboratory Animal Application and Research Center of Acıbadem Mehmet Ali Aydınlar University (ACUDEHAM). Each test was applied on different days and there was a day off between each test. Mice were carried to the testing room 1-hour before the experiment for their orientation to the environment. The arena was cleaned with 70% ethanol after each session of behavioural tests in order to eliminate olfactory cues. All of the experiments were recorded by a camera on the ceiling and analysed via Ethovision XT (Noldus Information Technology, Netherlands) system.

#### 4.5.1 Open Field Test

In order to assess locomotor activity, mice were placed in a 50×50 cm open field maze and each mouse was allowed to freely wander in the arena for 5 minutes. The open field arena was divided into 16 equal frames. The number of frames crossed by the mouse were recorded (***Seibenhener and Wooten, 2015***).

#### 4.5.2 Elevated Plus Maze Test

Anxiety was evaluated in the elevated plus maze apparatus. Mice were placed in the middle of a plus-shaped labyrinth with two open (50 × 10 cm) and two closed (50 × 10 × 40 cm) arms connected by a central square (10 × 10 cm) and elevated to a height of 50 cm. The time spent in the open and closed arms was noted separately for 5 minutes (***Komada et al., 2008***).

#### 4.5.3 Morris Water Maze

Spatial learning and memory were assessed by Morris Water Maze. The inside of the tank was filled with 24-26ºC water and the platform placed in a quadrant was kept 1.5 cm below the water level. Two sessions were carried out every day for four days. In each session, the mice were released into the water from four specific points each time and were allowed to swim for ninety seconds to find the platform. If they could not find it, the mice were placed on the platform for thirty seconds to give the opportunity to see environmental cues. The latency to find the platform was recorded and daily averages were calculated for each mouse. To test memory, on the fifth day, the platform was removed, and the mice were allowed to swim freely for ninety seconds once. Time spent in the quadrant that previously had the platform and the number of platform crossings were recorded (***Vorhees and Williams, 2006***).

### 4.6 Histopathological analyses

After the hippocampus samples were fixed with 10% neutral formalin, they were dehydrated by passing through 70%, 90%, 96% and 100% alcohol series. It became transparent with xylol and then embedded in paraffin (P3558-1kg Sigma Aldrich, Paraplast Embedding Media, U.S.A). Tissue sections are taken from paraffin blocks with a thickness of 5 μm. Then, they were stained with hematoxylin-eosin. These sections were examined under a light microscope (Zeiss) and evaluated in terms of inflammation, necrosis, tissue integrity and structural changes in cellular elements and apoptosis.

### 4.7. Western blot

The tissue samples were initially homogenized in the ice-cold RIPA lysis buffer (50 mM Tris–HCl, pH 7.4, 150 mM NaCl, 1% NP-40, 0.25 % sodium deoxycholate, 1 mM EDTA, 1 mM PMSF, 1 mM sodium orthovanadate, 1 mM sodium fluoride with 1X protease inhibitor cocktail) using tissue dounce homogenizer. The homogenized samples were then sonicated by Omni Ruptor 4000 sonicator approximately for 1 s. The lysates were centrifuged at 14,000Xg for 15 min. at 4°C to collect the supernatants. Concentration of extracted protein was quantified using Bradford protein assay (Bio-Rad, USA). Protein samples were separated by 10% SDS-PAGE (sodium-dodecyl sulphate-polyacrylamide gel electrophoresis) and then transferred to a nitrocellulose membrane using a semi-dry transfer system (40 min, 25 V) (Bio-rad, Transblot). The membranes were incubated with 5% bovine serum albumin for 1 hour at RT. Afterwards, the membranes were incubated with primary antibodies against to c-Jun (1:4000, 60A8, Rabbit mAb, Cell Signaling Technology), and GAPDH (1:2000) overnight at 4°C. The membrane was rinsed three times with PBS-T for 10 min., and then probed with appropriate secondary antibodies (1:5000) for 1 hour at RT and then washed three times with PBS-T for 10 min. The protein bands were visualized by an enhanced chemiluminescence reagent (SuperSignal™ West Femto Maximum Sensitivity Substrate, Thermo Scientific, USA). Finally, the relative densities of the protein bands were analyzed by Image Lab software (Bio-Rad, USA) (***Taylor and Posch, 2014***).

### 4.8 Statistical Analysis

Statistical tests were performed using Graph pad Prism 9.2.0 software (GraphPad Software Inc.). All data are presented as means ± SEM. Statistical significance was assessed by One-way ANOVA with post hoc Tukey-Kramer honest significant difference test for comparison across multiple groups. P value equal or less than 0.05 (p≤0.05) was considered significant.

## Acknowledgments

We thank Dr. Meltem Kolgazi for her contributions to biochemical analyses.

